# A randomized comparison of different hormone replacement protocols for thawed blastocyst transfer

**DOI:** 10.1101/114538

**Authors:** Sagiri Taguchi, Miyako Funabiki, Yoshihiro Tada, Masako Karita, Terumi Hayashi, Yoshitaka Nakamura

**Affiliations:** All authors address: Oak Clinic, Japan 2-7-9 Tamade-Nishi, Nishinari-ku, Osaka, 557-0045

## Abstract

**Background:** There are no randomized controlled trials evaluating the pregnancy rates after thawed blastocyst transfers in patients treated with various hormone replacement regimens.

**Methods:** A prospective randomized controlled trial was conducted to evaluate the outcomes in three different hormone replacement protocols for thawed blastocyst transfer. A total of 330 women (median age 38.2 years) who were undergoing IVF at our clinic were enrolled.

**Results:** Serum estradiol (E2) levels were 267.71 pg/ml in Premarin group, 391.22 pg/ml in Estrogel group and 495.12 pg/ml in Estrana tape group. Therefore, serum E2 levels in Estrana tape group were higher than those of the other two groups (P<0.01). The pregnancy rate in the Estrogel group was higher than that in the Premarin group (30.0% versus 17.3%, P=0.026, odds ratio 2.05, 95% confidence interval: 1.09–3.87). Furthermore, the pregnancy rate in the Estrana tape group was higher than that in the Estrogel group (43.6% versus 30.0%, P=0.036, odds ratio 1.81, 95% confidence interval: 1.04–3.14).

**Conclusion:** The serum E2 levels contributed to differences observed in the pregnancy rate among the three different protocols. Thus, Estrana tape has an advantage as a hormone replacement protocol for thawed blastocyst transfer.

## Introduction

The first successful pregnancy after use of frozen-thawed embryo transfer was reported in 1983 ^1^. Frozen-thawed embryo transfer is known as useful technology to increase cumulative pregnancy rate ^2^ Furthermore, frozen-thawed embryo transfer has accounted for around 20% of all assisted reproductive technology cycles in Europe in 2006 ^3^. However, as far as we know, there are no randomized controlled trials evaluating the pregnancy rates after thawed blastocyst transfers in patients treated with various hormone replacement regimens. Therefore, we investigated the clinical outcomes of three different hormone replacement protocols used in thawed blastocyst transfer procedures by a prospective randomized controlled trial.

## Methods

### Study design

A prospective randomized controlled trial was conducted to evaluate the outcomes in three different hormone replacement protocols for thawed blastocyst transfer. A total of 330 women (median age 38.2 years) who were undergoing IVF at our clinic were enrolled. The trial registration number was UMIN000016919. Participants were separated into three groups via computer-generated randomization.

### Regimens

Each regimens were as follows.

Premarin® group (n=110; 2.49 mg/day; from the second day of menstruation to the fifth day and 4.98 mg/day after the sixth day). Estrogel® group (n=110; 2.16 mg/day in Estrogen conversion after the second day of menstruation. Estrana tape® group (n=110; 2.88 mg, 4 pieces per 2 days).

### Institutional Review Board (IRB) approval

Our study was approved by the IRB of Oak Clinic, Japan. The patients provided informed consent at our clinic.

### Statistical tests

The statistical tests were performed using Dr. SPSS II for Windows (SPSS Japan, Inc., Tokyo), and significance was defined as p < 0.05 (two-tailed). Statistical analyses were investigated using univariate regression. Statistical significance was defined as p<0.05.

## Results

### Result 1

On the 12th day of the cycle, endometrium thickness was 9.19±0.31mm (mean±SD) in Premarin® group, 7.43±0.10mm (mean±SD) in Estrogel® group and 7.73±0.22mm (mean±SD) in Estrana tape® group.

Premarin® group versus Estrogel® group (P<0.001) or Estrana tape® group (P<0.001). Therefore, endometrium thickness in the Premarin® group were higher than Estrogel® group and Estrana tape® group.

### Result 2

Serum estradiol (E2) levels were 267.71±17.11 pg/ml in Premarin® group, 391.22±11.21 pg/ml in Estrogel® group and 495.12±18.54 pg/ml in Estrana tape® group. Therefore, serum E2 levels in Estrana tape® group were higher than those of the other two groups (P<0.01).

### Result 3

The pregnancy rate in the Estrogel group® was higher than that in the Premarin group® (30.0% versus 17.3%, P=0.026, odds ratio 2.05, 95% confidence interval: 1.09–3.87). Furthermore, the pregnancy rate in the Estrana tape® group was higher than that in the Estrogel group® (43.6% versus 30.0%, P=0.036, odds ratio 1.81, 95% confidence interval: 1.04–3.14).

### Result 4: Safety issues

The three different protocols were well tolerated, with no difference in the number of serious adverse events in the present study. Typical adverse drug reactions (breast discomfort and itching, *etc*) were observed in the three different protocols.

## Discussion

According to our prospective randomized controlled trial, the serum E2 levels contributed to differences observed in the pregnancy rate among the three different protocols. Thus, Estrana tape® has an advantage as a hormone replacement protocol for thawed blastocyst transfer. However, our study is limited to the three hormone replacement regimens studied for the patient population examined in the present study. In this regard, other hormone replacement regimens will be studied in the near future.

## Author contributions

All authors: Concept, Provision of the study materials, collection and/or assembly of the data, analysis and interpretation of the data and final approval of the manuscript.

## Acknowledgements

We are very grateful to nurses and clinical embryologists for their assistance with the design of this study and/or the experiments performed at our clinic.

